# Canopy micrometeorology monitoring in diverse climates innovatively improves the evaluation heat-induced sterility of rice under climate change

**DOI:** 10.1101/2020.11.01.363143

**Authors:** Mayumi Yoshimoto, Minehiko Fukuoka, Yasuhiro Tsujimoto, Tsutomu Matsui, Kazuhiro Kobayasi, Kazuki Saito, Pepijn A.J. van Oort, Baba I.Y Inusah, Chenniappan Vijayalakshmi, Dhashnamurthi Vijayalakshmi, W.M.W. Weerakoon, Silva Chandradasa, Tin Tin Myint Daw, Zar Chi Phyo Daw, Xiaohai Tian, Huu-Sheng Lur, Chwen-Ming Yang, Lee Tarpley, Norvie L. Manigbas, Toshihiro Hasegawa

## Abstract

This study is the first and unique of open-field studies to assess the determination process of heat-induced spikelet sterility (HISS) of rice by using the same variety and the same monitoring system, MINCER (Micrometeorological Instrument for Near Canopy Environment of Rice), covering the major-rice growing regions from Sub-Saharan Africa, South, Southeast, and East Asia, and USA. Applying the observation data from the monitoring network, MINCERnet, to the canopy heat balance model, IM^2^PACT, it was quantitatively corroborated in open-field conditions worldwide that the canopy and panicle transpiration and their evaporative cooling effect played a great role on the micrometeorological gap between the ambient air temperature and the panicle temperature, and that the sterility rate due to HISS in open-fields can be evaluated accurately in diverse climates by the mean panicle temperature at flowering hours in the flowering period. The heat balance structure suggested that the risk of HISS should be higher in high humidity climates rather than in dry climates also in the future, which lead to the importance of the humidity accuracy as well as of the air temperature in climate scenarios and their spatial downscaling for future prediction of rice heat stress and production. Applying the heat-tolerant variety was suggested to be able to keep the sterility due to HISS low in all climates. It is needed of the approach using the panicle temperature as indicator of HISS by intervening sub-model and/or monitoring of micrometeorology inside the canopy to reduce uncertainties in future rice yield prediction under various adaptation measures.

## 1 INTRODUCTION

In its 5th assessment report, the IPCC reported that the annual mean surface air temperature is most likely to increase by 0.3 - 4.8°C by 2100 depending on various GCMs under four RCPs (IPCC, 2013), and for the major crops in tropical and temperate regions, the climate change without adaptation will negatively impact on the production for local temperature increases of 2 °C or more above late-20th-century levels (IPCC, 2014).

Rice, the staple food for about three billion people, is well adapted to a rage of climates. It is an established fact, however, that rice is also highly susceptible to heat especially at flowering stage (Satake & Yoshida, 1978). Previous studies by growth chamber experiments revealed that the panicles become sterile when exposed to high temperatures over 35 °C for a few hours during flowering by poor anther dehiscence and low numbers of germinating pollen grains on the stigma, and that this heat-induced spikelet sterility (HISS) can be the major reason for yield losses (Matsui et al., 1999; Matsui et al., 2001; Prasad et al., 2006). Horie (1993) has developed an equation of the spikelet fertility percentage (100 – sterility percentage) as a function of daily maximum air temperature averaged over flowering period first in the world, from the TGC experiments. Since then, the daily maximum air temperature had been often referred to as a criterion for inducing the HISS. Researches have been conducted using closed system experiments such as growth chambers and TGC to investigate the relationship between the sterility and temperatures of the air/panicles to model them (e.g. Jagadish et al., 2007; Weerakoon et al., 2008). However, all these experiments evaluated the sterility rate at the scale of each panicle under controlled environments, and the obtained relational equations cannot be directly applied to the field, because of the temporal and spatial scale gaps between close system and open field.

One of the biggest gaps to induce the difficulty in evaluating the HISS in the field is the micrometeorological gap between above and inside the canopy. Matsui et al. (2007) demonstrated that the panicle temperature can be substantially lower than the air temperature above the canopy by as much as 6 to 7 °C under hot and dry conditions in the Riverina region of New South Wales, a major rice production area in Australia. On the other hand, Tian et al. (2010) reported that the panicle temperature can exceed air temperature by 4 °C under humid conditions, in the rice field of Jianghan Basin, China. One main reason for micrometeorological gap is the difference in the evaporative cooling effect due to canopy and panicle transpiration. That is, in Australia the panicle temperature decreased below the air temperature because of large transpiration stimulated by extremely dry and windy condition, while in China, the panicle temperature increased because the transpiration was suppressed by humid and low wind speed condition.

In recent years, some researches have used the panicle temperature or the thermal environment generated by plants themselves as indicators of HISS in the field condition instead of the ambient air temperature above the canopy. Julia and Dingkuhn (2013) estimated the panicle temperature in the field empirically from weather station data, to which the observed sterility rate could be related for contrasting climates. van Oort et al. (2014) proposed a simple model of the HISS as a function of flowering time distribution and the panicle temperature in the field, where the panicle temperature was simply estimated using the relative humidity since it is the main factor to drive the plant transpiration and its cooling effect. These studies showed the need to consider the canopy microclimate and organ temperature for reducing the uncertainties in predicting the global warming effect and the adaptation study. Yoshimoto et al. (2011) has developed the micrometeorology model to estimate the panicle temperature in rice canopies, IM^2^PACT. Such a modeling approach is effective to estimate the panicle temperature and the canopy micrometeorology directly from the heat balance process, but meteorological observation data with sufficiently guaranteed accuracy is required not only as input data to the model but also as model validation data.

Observational data of the thermal environment inside the canopy with the accuracy enough for climate change impact study was lacking or completely missing so far, in agricultural fields in the world. One of the reasons is the difficulty of measuring the air temperature inside the canopy in the field, which generally requires a ventilation shield to minimize the effects of solar radiation. Force-ventilation is the best option for accuracy, but it requires a certain amount of electric power to drive the fan, which is difficult in paddy field without power supply. On the other hand, the natural ventilation shield does not require a power source for ventilation, but the air temperature cannot be measured correctly inside the canopy because the natural ventilation is obstructed by vegetation. Therefore, in order to bridge the micrometeorological gap above and inside the canopy with sufficiently accurate observation data and reduce the uncertainty regarding the evaluation of HISS in the field, we have developed an instrument MINCER, which can measure the air temperature and the humidity inside the canopy by force-ventilation without external power supply (Fukuoka et al., 2012a). And we have developed a multi-site monitoring network MINCERnet by deploying MINCERs to ‘hot spots’ of rice producing areas in the world’s various climates.

The objectives of this paper are to clarify (1) the actual state and the heat balance mechanism of micrometeorological gap above and inside the canopy in a wide range of climates with observation data from MINCERnet, 2) the relationship between the HISS observed in the field and the panicle temperature at flowering hours estimated by the model IM^2^PACT, and 3) the risk of HISS in the monitoring sites under the current climate. We hypothesized that the approach using the panicle temperature or the thermal environment inside the canopy as indicators of HISS should be effective and indispensable not only for impact assessment but also for adaptation studies.

## 2 MATERIALS AND METHODS

### 2.1 Experimental design in the monitoring network

Sites and their description of MINCERnet were shown in Figure 1 and Table 1. The initial network had been deployed in eight sites of the world’s rice growing regions, India, Sri Lanka, Myanmar, the Philippines, China, Taiwan, Japan and the USA sites in 2010 (Yoshimoto et al., 2012), where the preliminary monitoring of thermal environments had been conducted using the respective standard variety different among sites. The target seasons was the hottest season and the moderately hot season in a year. We included Senegal, Benin and Ghana in Sub-Saharan Africa since 2014. Then the monitoring network has been formed consisting of totally eleven sites as of 2017 shown in Figure 1.

**Figure 1.**
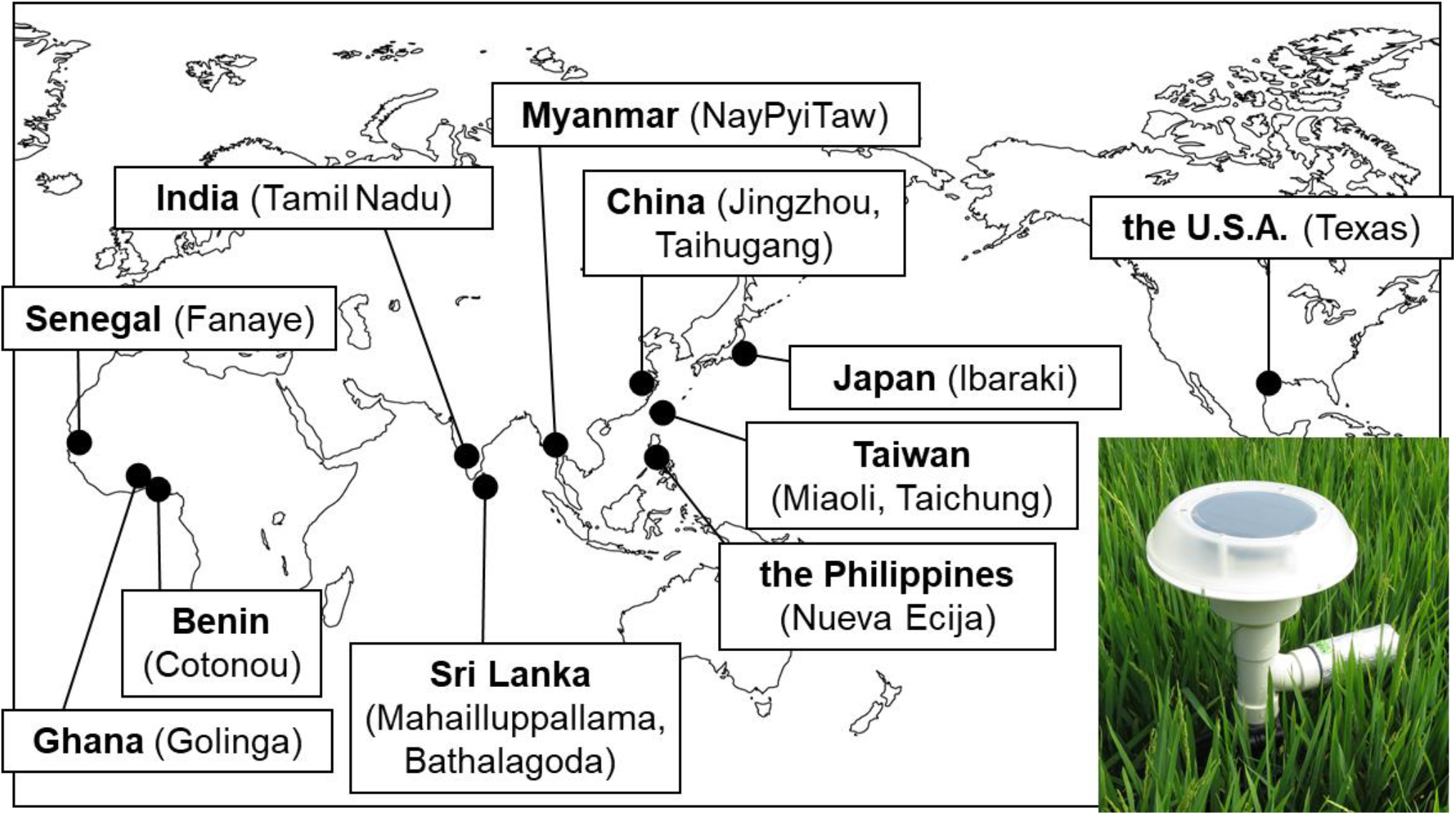
Sites of the multi-site monitoring network, MINCERnet as of 2017, and the MINCER instrument.

**Table 1.** Geographic information and experimental year of MINCERnet sites

| Symbols | Site |  | Latitude | Longitude | Elevation<br>(m) | Experimental year |
| --- | --- | --- | --- | --- | --- | --- |
| SN | Senegal | Fanaye | 16.54 | -15.19 | 9 | 2014-present |
| GH | Ghana | Golinga | 9.35 | -0.95 | 108 | 2016-present |
| BJ | Benin | Cotonou | 6.37 | 2.39 | 3 | 2014-2015 |
| IN | India | Tamil Nadu | 11.00 | 76.92 | 422 | 2010-2013, 2017-present |
| LK | Sri Lanka | Mahailluppallama | 8.10 | 80.47 | 122 | 2010-2019 |
|  |  | Bathalagoda | 7.53 | 80.44 | 123 | 2019-present |
| MM | Myanmar | NayPyiTaw | 19.82 | 96.27 | 96 | 2010-2011, 2017-present |
| PH | Philippines | Nueva Ecija | 15.67 | 120.89 | 58 | 2013-present |
| CN | China | Jingzhou | 30.35 | 112.15 | 34 | 2010-2014 |
|  |  | Taihugang | 30.36 | 112.05 | 30 | 2017-present |
| TW | Taiwan | Miaoli | 24.48 | 120.82 | 116 | 2010-2013 |
|  |  | Taichung | 24.03 | 120.70 | 80 | 2016-present |
| JP | Japan | Ibaraki | 36.03 | 140.11 | 22 | 2010-present |
| US | USA | Texas | 30.07 | -94.29 | 10 | 2010-2013, 2017-present |

Since 2017, the target of MINCERnet was shifted to quantifying the relationship between the HISS and the thermal environment over a wide range of climates by using a common variety, IR64, the moderately susceptible variety against the HISS (Jagadish et al., 2010) for all sites. In addition to the common standard variety of IR64, the heat tolerant variety, N22 (Jagadish et al., 2010; Mackill et al., 1982) was cultivated in Senegal, India, Sri Lanka, Myanmar, the Philippines, China and Japan sites. And the heat susceptible variety, IR52 (Mackill et al., 1982) was cultivated in India, the Philippines and Japan sites. In the tropics in Senegal, Ghana, India, Sri Lanka, Myanmar and the Philippines sites, both dry and wet seasons were targeted to clarify the characteristics of the thermal environment when the weather conditions are significantly different in a year.

Field cultivation trials have been conducted at flooded rice paddies by conventional management in each study region. Three replicated plots were provided, with each plot large enough to maintain a uniform microclimate inside the rice canopy. A randomized complete block design with three replicated plots was used in trials using multiple varieties since 2017. An area for installing MINCER was provided near the center of the plot to monitor the air temperature and relative humidity inside the canopy. In addition, the drainage treatment plots were also set up at the same time to investigate the effects of drought stress in Ghana, Sri Lanka, Myanmar, the Philippines, Taiwan and the USA sites (Tsujimoto et al., 2021 for Ghana). At these sites, the data of the flooded plots as the control plots were analyzed in this study.

### 2.2 Micrometeorology measurement above and inside the canopy

MINCER (Micrometeorological Instrument for Near Canopy Environment of Rice) is a stand-alone force-ventilated measurement system (inserted in Figure 1) which was developed by Fukuoka et al. (2012) for the current monitoring network, MINCERnet. MINCER uses a solar cell powered ventilator equipped with rechargeable batteries (Solar Vent MPV; ICP Solar Technologies Inc., Canada) and a power control circuit proprietarily developed for MINCER. An air temperature and humidity sensor with logging function (RX-350TH, As One Corp., Japan) is housed inside the ventilated radiation shield. The system is lightweight and can be settled at any desired height in the rice canopy using an adjustable tripod. MINCER can be easily handled by agronomists and crop physiologists, even if they were not experts in agricultural meteorology, and ensure that anyone can conduct accurate measurements of the air temperature in the field without power supplies.

MINCERs were installed above and inside the canopy in the field targeting the heading period and later. MINCERs for inside-canopy measurement were set in one or more replicated plots for all varieties in the site, and were installed at the height of the panicles between rows to monitor the air temperature and the relative humidity inside the canopy. And one more MINCER was installed in the same field at a height set to twice the canopy height or higher, to monitor the air temperature and the relative humidity above the canopy. The air temperature and humidity by MINCER were recorded at 2-minutes intervals 24 hours per day.

If there was a nearby weather station with hourly or finer time intervals data available in the sites, the air temperature and humidity there were also incorporated into the analysis with those by MINCERs. The solar radiation and the wind speed at the nearby weather station were used for the heat balance calculation to estimate the panicle temperature. At the sites where the solar radiation and wind speed data at the weather station were not available, an additional simple system for measuring them was provided in the field. The system included a pyranometer (CMP3, Kipp and Zonen B.V., the Netherlands) and the weather transmitter (WXT520, Vaisala Oyj, Finland), by which the solar radiation and the wind speed were measured every 10 seconds and recorded their 10-minutes average values by the datalogger (CR850, Campbell Sci. Inc., USA) powered by the solar panel (SP10, Campbell Sci. Inc., USA).

### 2.3 Crop monitoring and sterility measurement

Each replicated plot had a heading monitoring area and a flowering monitoring area of about 1m x 1m. In the heading monitoring area, the number of emerged panicles were counted. The state in which the tip of the panicle emerges from the leaf sheath is defined as ‘heading’, and the day when the number of headed panicles reaches 50% of the total number of panicles is defined as the ‘heading day’. In the heading monitoring area, in addition to this 50% heading date, the days when heading started and when all panicles headed were recorded.

In the flowering monitoring area, flowering (spikelets opening) time in a day, FOT, was observed by overviewing. As there was a concern about bias in the overviewing depending on the observers, a time-lapse photography with a digital camera was introduced for objective identification since 2017. The canopy surface with emerged panicles was photographed at 10-minutes intervals with waterproof digital cameras (WG-40/WG-60, RICOH Imaging Company Ltd., Japan) for a several days in the flowering period. The digital cameras were fixed above the canopy and inclined 45 degrees from the horizon in the most representative plot for each variety. The times on FOT were identified from the captured images. The ‘initial’ flowering time was defined as the time when at least one spikelet opened visible in the photographed area. The ‘peak’ flowering time was the median of the distribution of spikelets opening time, that is, the time when the maximum number of spikelets had newly opened during the 10 minutes.

All the panicles of four hills surrounding the intake of MINCERs were sampled and threshed for investigating the spikelet sterility rate. The sterility rate was determined by counting the numbers of sinking and floating spikelets in 70 % ethanol, as per Kobata et al. (2010).

### 2.4 Panicle temperature estimation

The canopy surface temperature and the panicle temperature have been manually measured with a high-accuracy handheld infrared radiometer (MI-230, Apogee Instruments, Inc., USA) since 2017. The measurement was conducted once a day in flowering hours for 3 days in the flowering period and around noon to the hottest hours for 3 days with about one-week interval in maturing period. One measurement included two readings of the canopy surface temperature and three readings of the panicle temperature for each replicated plot.

The canopy surface temperature was measured by setting the MI-230 sensor at >50 cm above the canopy surface with an angle of 45 degrees. In this way, the average temperature of an elliptical area of about > 1500 cm^2^ was measured. The panicle temperature was measured by collecting 2 to 3 panicles intact in the field and holding them lightly to make ‘panicle wall’ so that the field of view of the radiometer should not include the background behind the panicles.

The IM^2^PACT (Yoshimoto et al., 2011) is the physical heat balance model of bulk canopy to estimate the panicle temperature originally targeting Japanese rice varieties in rice paddy field in Japan. The IM^2^PACT, briefly, solves the heat balance between the air above the canopy and the bulk canopy first, to calculate the bulk canopy temperature and to estimate the air temperature and humidity inside the canopy, and solves the heat balance between the air inside canopy and the panicle in it second, to calculate the panicle temperature, T_p_. The heat balance calculations in the IM^2^PACT requires hourly data of meteorology such as the solar radiation and wind speed as well as the air temperature and humidity above the canopy, and the vegetation parameters must be set such as the canopy height, leaf area index and the transpiration intensity of canopy and panicle. One of the important parameters that directly affect the plant body temperatures is the transpiration conductance of the body surface. The parameters of the canopy and panicle transpiration conductance were determined so that the calculated values of the canopy and panicle temperatures match their measured values. The model was also validated by comparing the air temperature and the humidity inside the canopy, which is calculated as an intermediate output in the model, with their measured values by MINCER.

### 2.5 Data processing and statistical analysis

The quality check was conducted of all the air temperature and the relative humidity data by MINCER for all sites. In this study, the 2-minutes interval data by MINCER were averaged into hourly data to create as an hourly database with the weather station data, because the nearby weather station data was hourly in general. Drawing the box plot and the regression curve fitting were conducted with KaleidaGraph software 4.0.

## 3 RESULTS

### 3.1 Micrometeorological gap between above and inside the canopy

Figure 2 shows the example of distributions of the air temperature and the relative humidity in 24 hours of 29 days including the respective 50% heading day (days after heading, DAH, is −7 to +21) at all the experimental sites. The detail information for the Figure 2 are listed in Table 2, where the years/seasons were chosen from ones with the characteristic climate of the respective site to have all the measurement data as much as possible, at nearby weather station, above the canopy and inside the canopy.

**Figure 2.**
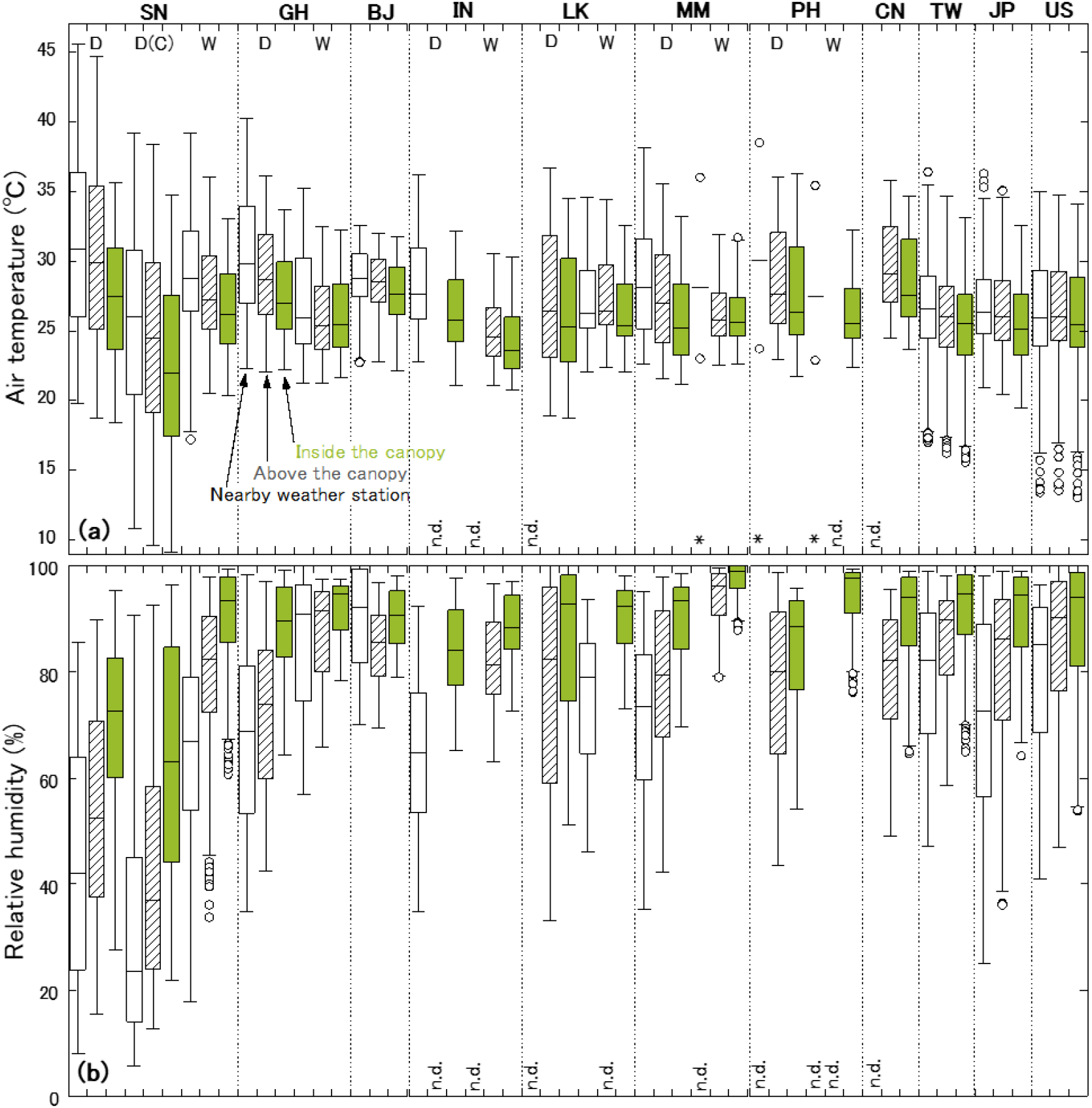
Distributions of (a) the air temperature and (b) the relative humidity at the nearby weather station, above the canopy and inside the canopy of 24 hours in the observation period, by box-plots. The “n.d.” means no data because of no available data from nearby weather station or sensor malfunction. At the three seasons in MM and PH marked by “*”, the daily maximum, mean and minimum air temperatures at the nearby weather station were plotted as an alternative of box-plots.

**Table 2.**
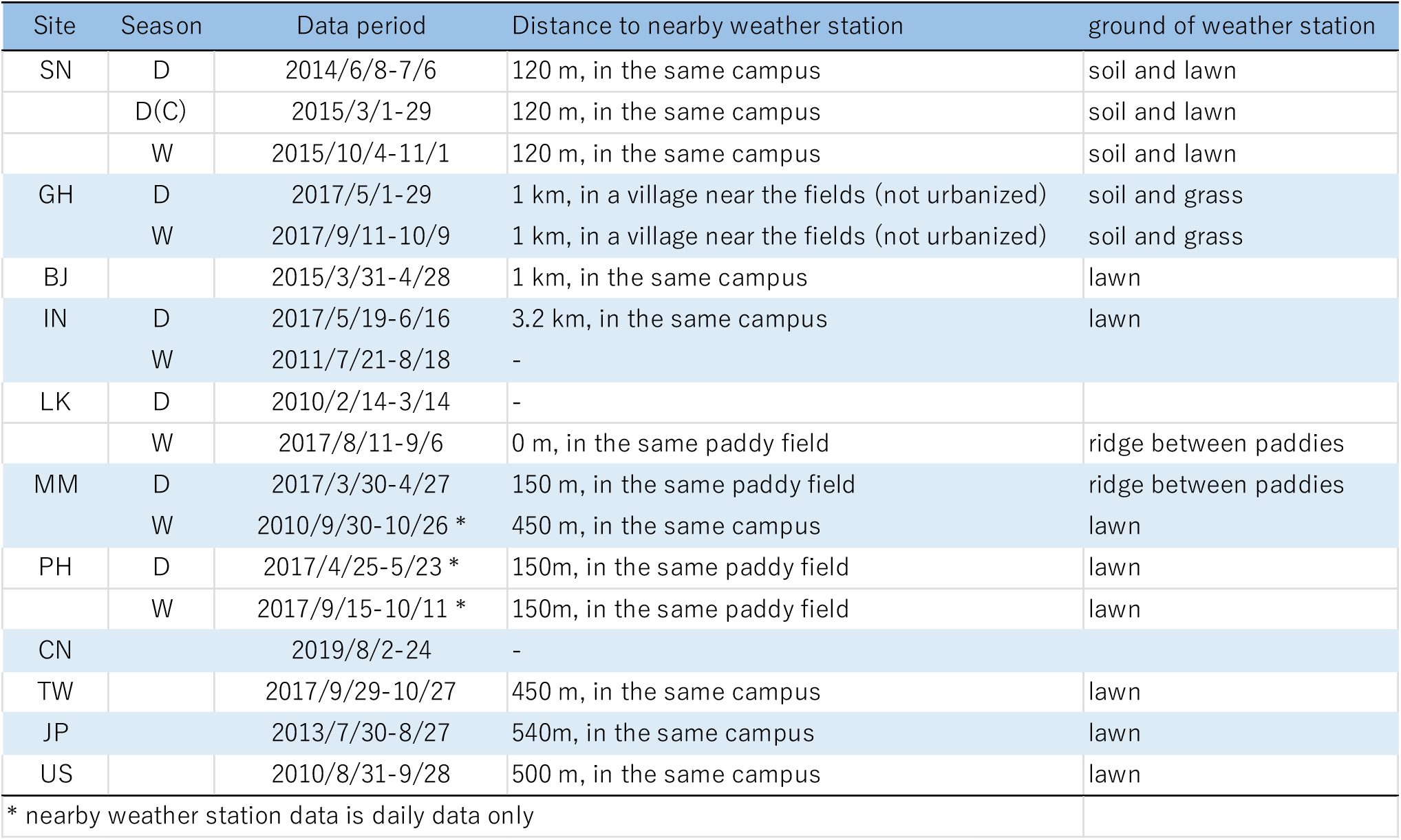
Observation period of each site chosen for Figure 2, and the information on the nearby weather station. The data period is 29 days at most including the respective heading day; days after heading, DAH is from –7 to +21.

| Site | Season | Data period | Distance to nearby weather station | ground of weather station |
| --- | --- | --- | --- | --- |
| SN | D | 2014/6/8-7/6 | 120 m, in the same campus | soil and lawn |
|  | D(C) | 2015/3/1-29 | 120 m, in the same campus | soil and lawn |
|  | W | 2015/10/4-11/1 | 120 m, in the same campus | soil and lawn |
| GH | D | 2017/5/1-29 | 1 km, in a village near the fields (not urbanized) | soil and grass |
|  | W | 2017/9/11-10/9 | 1 km, in a village near the fields (not urbanized) | soil and grass |
| BJ |  | 2015/3/31-4/28 | 1 km, in the same campus | lawn |
| IN | D | 2017/5/19-6/16 | 3.2 km, in the same campus | lawn |
|  | W | 2011/7/21-8/18 | - |  |
| LK | D | 2010/2/14-3/14 | - |  |
|  | W | 2017/8/11-9/6 | 0 m, in the same paddy field | ridge between paddies |
| MM | D | 2017/3/30-4/27 | 150 m, in the same paddy field | ridge between paddies |
|  | W | 2010/9/30-10/26 * | 450 m, in the same campus | lawn |
| PH | D | 2017/4/25-5/23 * | 150m, in the same paddy field | lawn |
|  | W | 2017/9/15-10/11 * | 150m, in the same paddy field | lawn |
| CN |  | 2019/8/2-24 | - |  |
| TW |  | 2017/9/29-10/27 | 450 m, in the same campus | lawn |
| JP |  | 2013/7/30-8/27 | 540m, in the same campus | lawn |
| US |  | 2010/8/31-9/28 | 500 m, in the same campus | lawn |
| * nearby weather station data is daily data only |  |  |  |  |

In tropical sites with dry (D) and wet (W) seasons, the humidity level was higher in the wet seasons than in the dry seasons. The air temperature level generally tended to be higher in the dry seasons than in the wet seasons.

The range of daily temperature variation was the largest in the dry seasons in Senegal site among sites. The range observed above the canopy was more than 20 °C. The daily change in temperature was extreme in Senegal site especially in the dry season as notated by D (C) in Figure 2, where the air temperature rose to nearly 40 °C in daytime, while it dropped to 10 °C or less in nighttime. The range of daily temperature variation was the smallest about 10 °C in Benin site among sites. It was small also in wet seasons in Ghana, India, Myanmar sites and in China site. In those sites, the nighttime temperature level was high, especially in China site.

Among the measurements at three positions; at nearby weather station, above the canopy and inside the canopy, the air temperature was the highest and the relative humidity was the lowest at nearby weather station. The air temperature became lower and the humidity higher as approaching to the plant canopy from at nearby weather station to inside the canopy. It is because of the transpiration of plant canopy and the evaporation of water surface, which decreases the canopy temperature by their evaporative cooling effect as well as increases the humidity adjacent to the canopy.

All the nearby weather stations were not located far from the experimental paddy field sites (Table 2) with the same altitude and are not urbanized. Despite it, Tst tended to be higher than Tab because the weather stations were typically not located in the paddy field. While in the wet season trial in Sri Lanka, T_st_ is almost same as T_ab_ because the nearby weather station was installed in the paddy field.

The air temperature inside the canopy, Tin tended to be lower than T_ab_ generally in all experimental sites. The temperature difference between inside and above the canopy (T_in_ – T_ab_) was highly related to the humidity level (Figure 3). This is because the canopy transpiration and thus its evaporative cooling effect is related to the humidity. T_in_ was lower than T_ab_ by 9.6 °C in the dry season of Senegal site because of its extremely low humidity level and strong evaporative cooling. In contrast, the magnitude of T_in_ – T_ab_ was much small, such as in the wet seasons in Ghana, Myanmar, Benin and India sites, and in the temperate sites. Under very high humidity condition in a day, T_in_ can be even higher than T_ab_ because the evaporative cooling effect is suppressed by low evapotranspiration.

**Figure 3.**
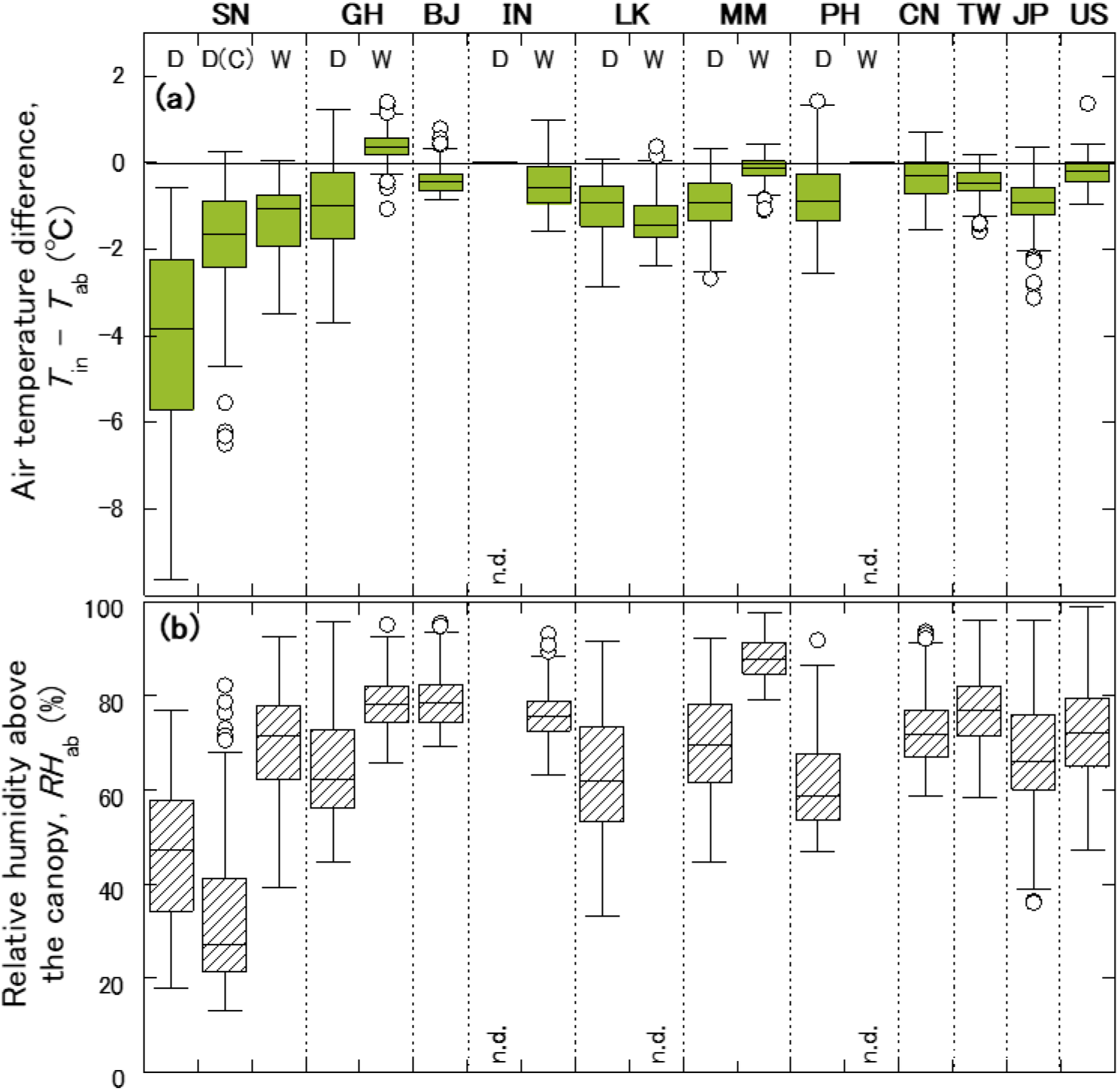
Distributions of (a) the air temperature difference between above and inside the canopy (T_in_ – T_ab_), and (b) the relative humidity above the canopy, RHab of daytime 6 hours (9:00-15:00 in local time) in the observation period, by box-plots. The “n.d.” means no data because of sensor malfunction.

### 3.2 Sterility and thermal variables

Sterility rate of all varieties obtained in all sites so far were investigated in the light of meteorological data. A high sterility rate probably due to low temperature was occasionally observed, often in the dry season in Senegal site as notated by D (C) in Figure 2 and rarely in other sites such as Sri Lanka and Japan sites. The sterility rate data of the experiment in which the T_st_ or T_ab_ was often well below 20 °C in the two weeks before the heading date had been excluded in this study focusing on heat induced spikelet sterility.

Figure 4 shows the relationship between the sterility rate of all varieties and thermal variables. The thermal variables were averaged for 7 days from 50% heading date (days after heading, DAH, is 0 to 6), as the actual main flowering (spikelets opening) occurs later than the 50% heading date and its peak is typically 2-3 days after the heading date. The daily maximum air temperature at weather station is one of the easiest information to obtain anywhere and often used to evaluate the heat stress risk. When applying the daily maximum of T_ab_ as the thermal variable, there was little correlation with sterility rate. The correlation coefficient, however, was slightly higher when applying the daytime (9:00-15:00 in local time) average of T_ab_ instead of its daily maximum. And the correlation was still higher when applying the daytime average of T_in_, where the data points of high sterility rate had the high T_in_. It seems certain that thermal variables closer to the plant canopy such as T_in_ can more accurately assess the sterility rate, although there is a still large variability since various varieties data were included.

**Figure 4.**
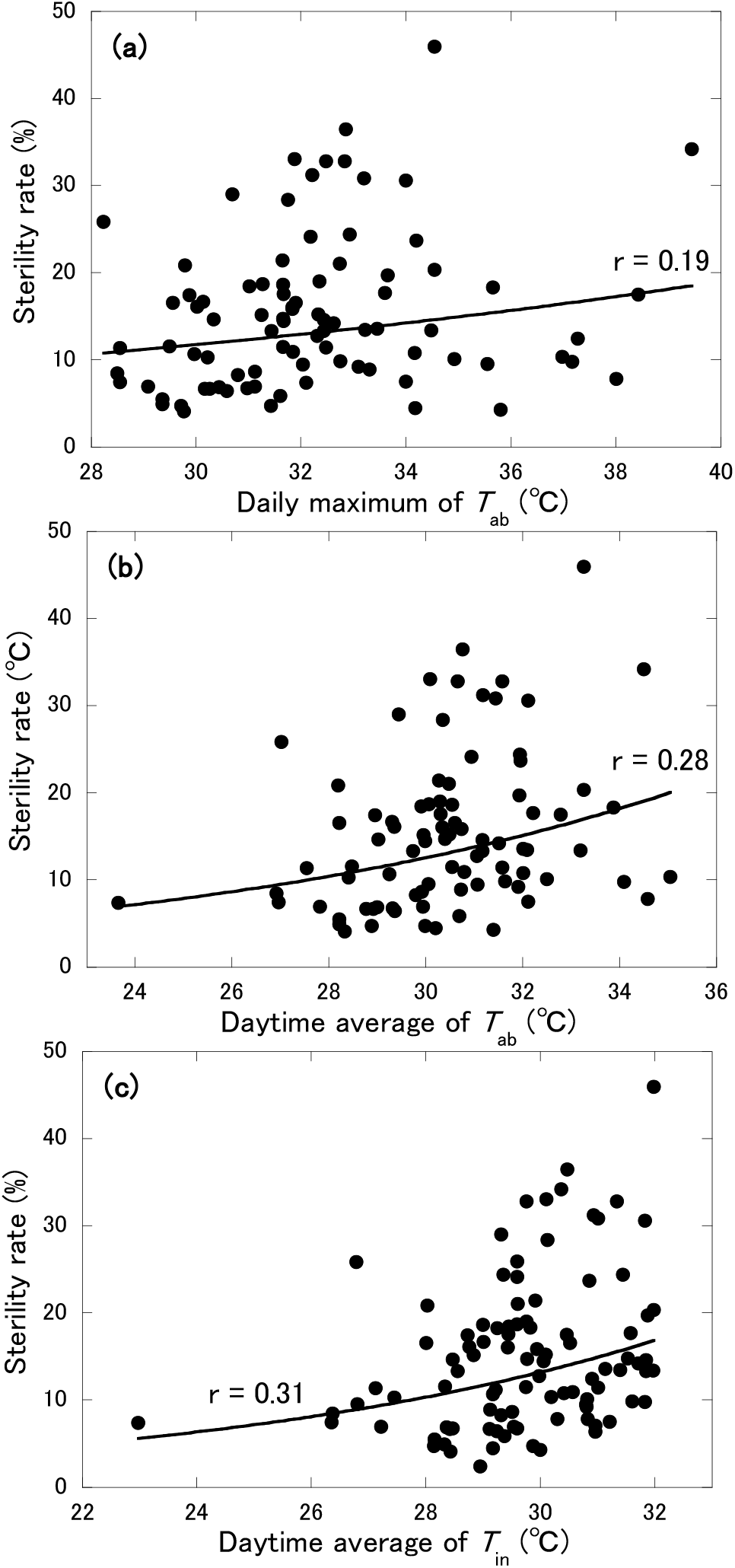
Relationship of the sterility rate of all varieties with thermal variables; (a) daily maximum T_ab_, (b) daytime average of T_ab_ and (c) daytime average of T_in_. All the thermal variables were averaged for 7 days from the respective heading day.

### 3.3 Sterility of the common varieties

#### 3.3.1 Flowering time in a day

To explore the thermal variables that determine the sterility rate, analysis was focused on the trials using IR64, N22 and IR52, eliminating variations due to different varieties. Flowering (spikelets opening) time in a day, FOT, is one of key factors relating to the heat induced spikelet sterility. To examine the relationship with the sterility rate roughly in Figure 4, it was attempted to apply the same time zone (6 hours of 9:00-15:00 in local time) to all sites as a daytime relating to flowering event, however the actual flowering time zone is not always the same among sites. Figure 5 shows the distributions of FOT objectively measured by time-lapse photography with a digital camera. Data in Benin site is exceptionally by overview information, because the experiments in Benin site were conducted before introducing objective measurements by time-lapse photography. The peak time of flowering of IR64 was averagely around 12:00 in Senegal, Benin and USA sites, 11:00 in Ghana site, and 10:00 in other sites, in each local time. The initial time of flowering was distributed averagely 1 hour ahead of the peak time.

**Figure 5.**
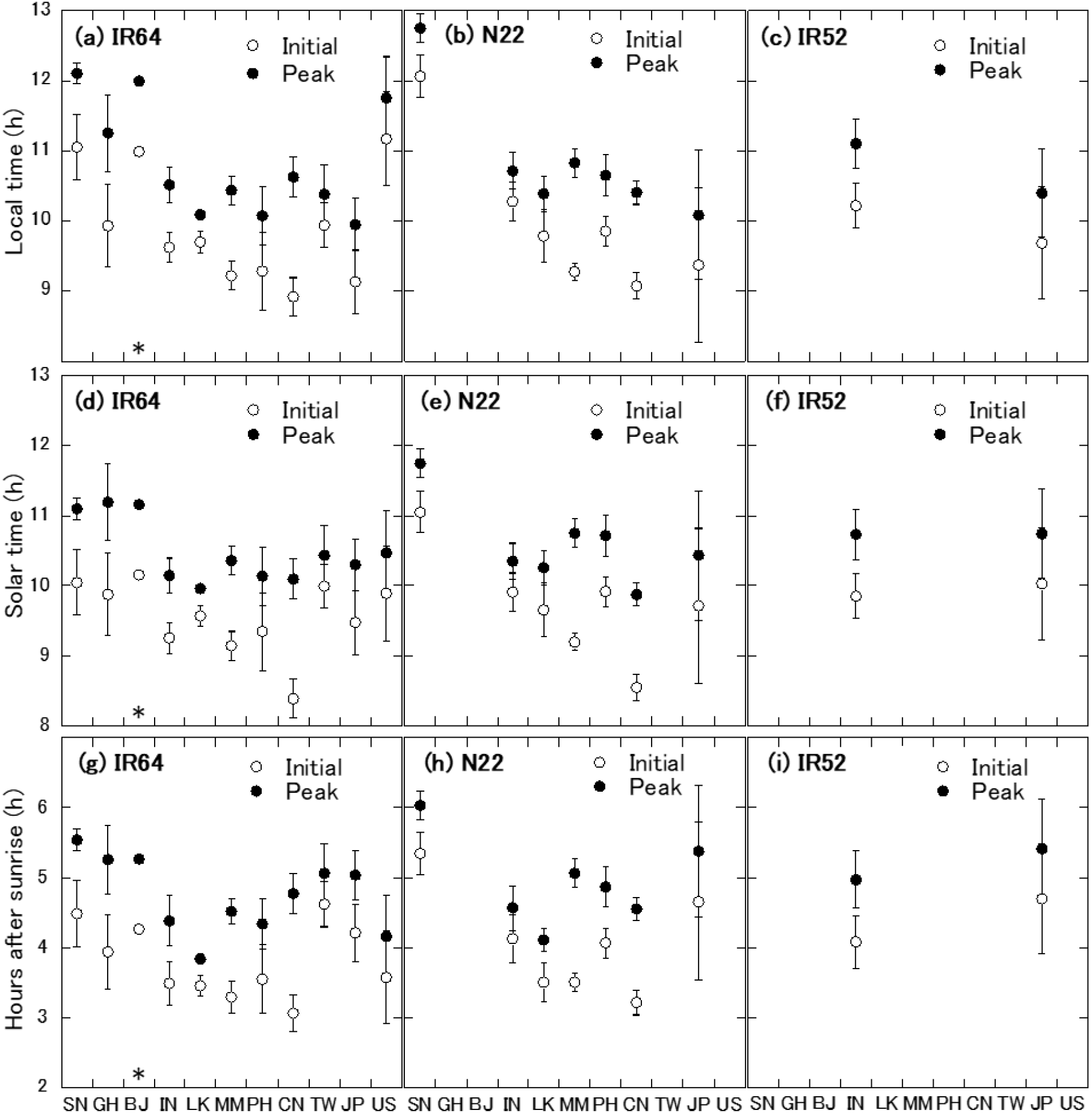
Distributions of flowering (spikelets opening) time in a day, FOT, of IR64, N22 and IR52, shown in local time (a-c), in solar time (d-f) and in hours after sunrise (g-i). The initial and peak times were objectively analyzed by time-lapse photography with a digital camera. Exceptionally in BJ (Benin site) marked by *, the values were by overviewing.

These observed FOTs in local time were converted to solar time and hours after sunrise (hasr) by solar geometry with the geographic information. Expressed as solar time, the variation in the peak time of flowering of IR64 among sites was reduced, around 11:00 in Ghana, Senegal, and Benin sites and 10:00 in other sites, however there was still a difference of FOT among sites despite the same variety. Expressed as hasr, there was also still a difference of FOT among sites. Those difference among sites may be because of environmental factors. In the field condition instead of controlled environments, various meteorological factors may affect FOT, which can affect the mean flowering time zones reflected by the characteristics of each climate. Kobayasi et al. (2010) revealed that the air temperature, solar radiation and vapor pressure deficit in early morning significantly affected FOT of a certain variety. Julia and Dingkuhn (2012) showed the possibility to explain the peak time of FOT by the mean minimum air temperature during the 7 days before anthesis across different varieties and environments. Incidentally, the seasons of FOT observed in Senegal and Philippines sites in this study are corresponded to Senegal hot-dry season and Philippines dry season in their study, where the peak time of FOT was distributed in the same range, around 4 to 6 hours in hasr, in both studies.

It is still unclear which environmental factors affected the FOT difference among sites. In this study, however, the flowering time zones in all sites were determined based on the actual distribution of FOT in local time instead of the same time zones of 9:00-15:00 among sites; as 6 hours of 10:00-16:00 in Senegal, Benin and USA sites, of 9:00-15:00 in Ghana site, and of 8:00-14:00 in the other rest sites, because the meteorological variables dataset is organized by local time. The peak time of flowering of N22 and IR52 tended to be slightly later than of IR64. The flowering 6 hours of N22 and IR52 were set as the same as of IR64 in the following analysis for convenience, because the meteorological variables dataset is hourly based, which is coarser time resolution than the difference of FOT among varieties.

#### 3.3.2 Panicle temperature

The panicle temperature, T_p_, was calculated with the physical heat balance model, IM^2^PACT, for all the experiments where hourly data of the solar radiation and wind speed were available at nearby weather station in addition to T_ab_ and RH_ab_ measurement by MINCER. As a result of determining the parameters of the canopy and panicle transpiration conductance so that the calculated values of canopy and panicle temperatures match the measured values at each site, it was found that their measured values cannot perfectly but can almost reproduced even if the common parameters for transpirations are applied across the sites, seasons and varieties.

To apply the IM^2^PACT to Indica rice varieties under diverse climatic conditions, however, three major modifications were needed from the original parameters for Japanese rice varieties under the climate in Japan, which are briefly as below: (1) The magnitude of transpiration conductance of IR64, N22 and IR52 were determined larger than of Japanese varieties, for both of canopy and panicle. (2) The dependence of panicle transpiration conductance on humidity was eliminated. With Japanese varieties and in a narrow range of high humidity environment in Japan, the reproducibility tended to increase with the parameterization that the panicle transpiration conductance decreased as the humidity decreased, but such parameterization failed in this study when extrapolating to the low humidity condition. Rather, if the transpiration conductance in daytime was constant regardless of humidity, the panicle temperature could be reproduced for Indica varieties under a wide range of climatic conditions. (3) The ratio of incident solar radiation on panicle to the global solar radiation was set smaller in N22 than in IR64 and IR52, because the canopy height was much higher and the panicles at flowering period were located relatively deeper inside the canopy in N22 compared to other varieties.

Figure 6 shows the comparison of estimated and measured T_p_ for three varieties in Senegal, China and Japan with contrasting climate conditions. Despite the simplification of the model parameterization, such as by applying common transpiration parameters among sites and varieties, or by eliminating its humidity dependence, the IM^2^PACT could almost reproduce the panicle temperature in daytime (RMSE=1.4 °C). The actual panicle temperature in the field can fluctuate every moment and have trend due to meteorological changes. While the estimated T_p_ is calculated by using the hourly averaged value based on continuous measurement, the measured T_p_ is by spot measurement for a several minutes in the hour, which was attempted to time average of the fluctuation and trend by replicated measurement though. Therefore, the both T_p_ values do not necessarily match perfectly. In Figure 6, however, whether T_p_ is high or low with respect to T_in_ is almost the same between the measured T_p_ and the estimated T_p_. The both T_p_ tended to be lower than T_in_ in Senegal site (dry season), where the relative humidity inside the canopy RH_in_ was lower than in other sites (Figure 2) and the T_p_ was likely to decrease by the large evaporative cooling effect due to larger panicle transpiration. While in China site, the RH_in_ was high and the T_p_ was likely to increase by the small evaporative cooling effect due to small panicle transpiration. In Japan site, the T_p_ was almost the same as T_in_, but sometimes higher or lower than T_in_ due to daily changes in weather conditions.

**Figure 6.**
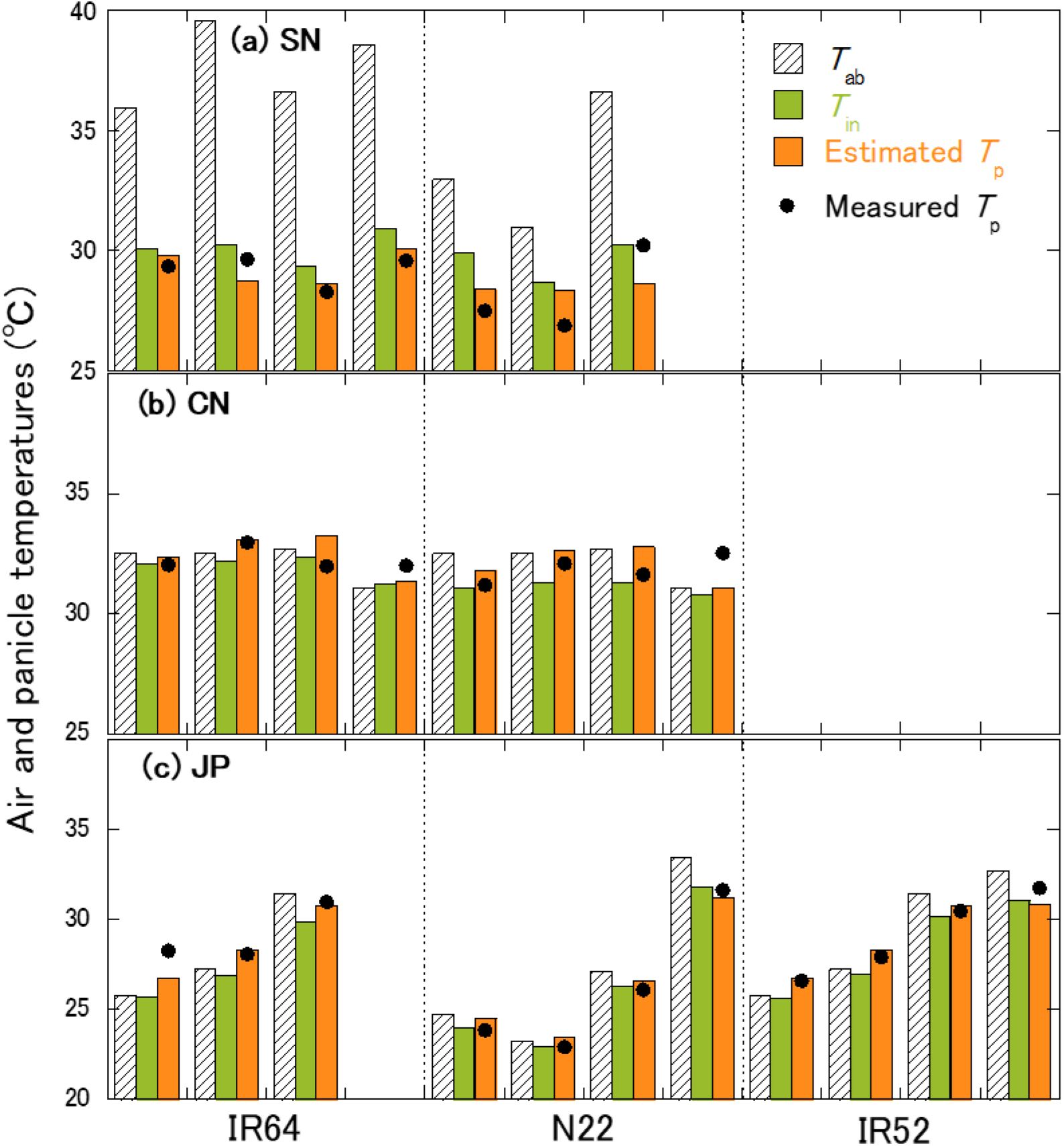
Comparison of estimated and measured T_p_ for IR64, N22 and IR52 in (a) Senegal, (b) China and (c) Japan sites contrasting climate conditions. Columns show the hourly data of measured air temperatures, T_ab_ and T_in_ and estimated panicle temperature, T_p_ by IM^2^PACT. Dots show the measured T_p_ by the infrared radiometer.

For the seasons in the sites shown in Figure 2, the T_p_ of 6 hours in the flowering time zones shown in 3.3.1 of 29 days period was calculated, and their distributions were shown together with the measured T_ab_ and T_in_ distributions in the same flowering time zones (Figure 7). Compared by median, the estimated T_p_ tended to be lower than T_in_ in the dry seasons of Senegal and Philippines sites, where the relative humidity inside the canopy, RH_in_ was low, while the estimated T_p_ tended to be higher than T_in_ in Ghana, Benin, and Philippines (wet season) sites and the temperate sites where RH_in_ was high. The panicle temperature is determined by the heat balance inside the canopy related to various meteorological factors such as solar radiation and wind speed, and is not determined only by the evaporative cooling effect that is largely related to RH_in_. It is certain, however, that the humidity environment inside the canopy created by the plant canopy via its transpiration has a great influence on the determination process of the T_p_.

**Figure 7.**
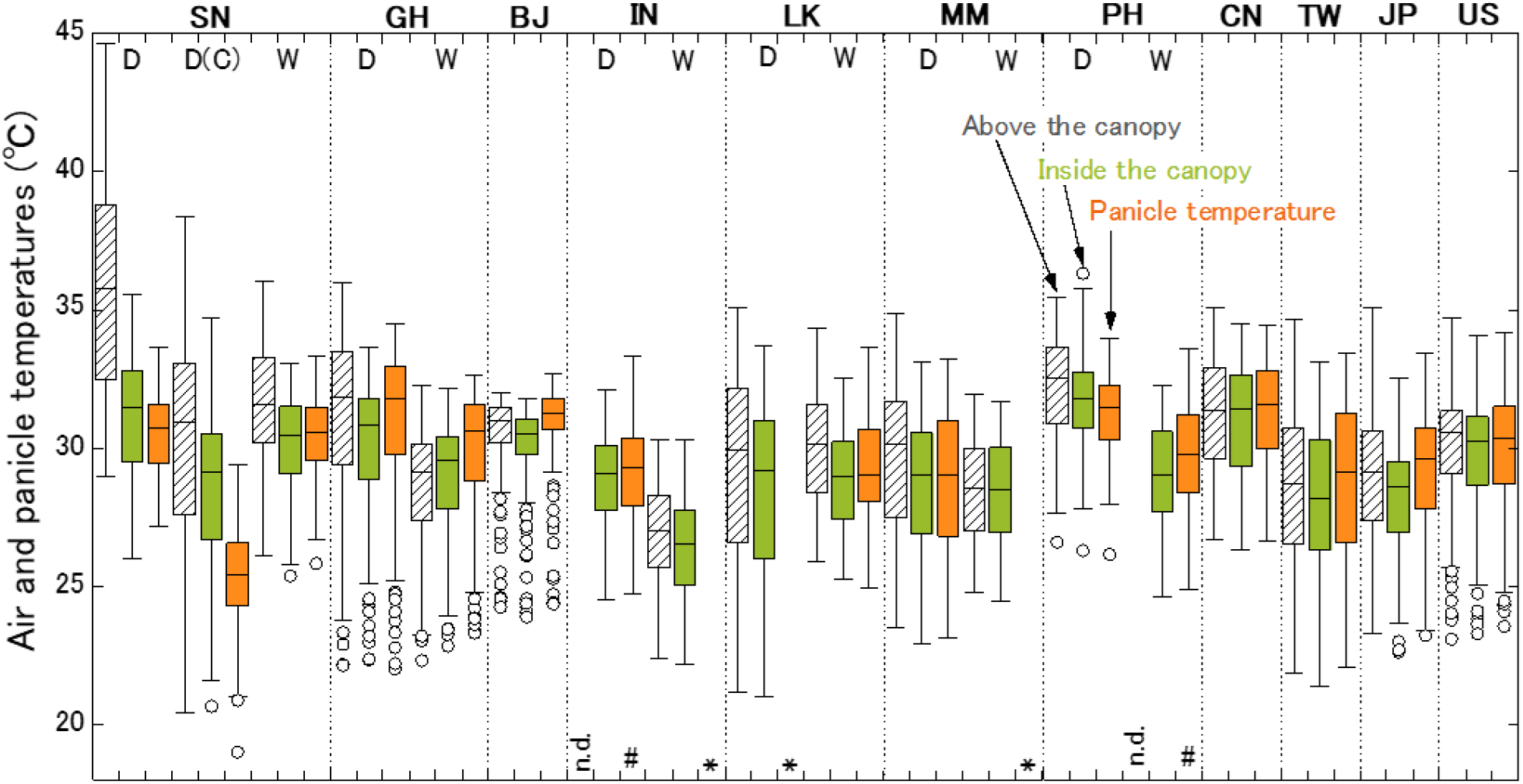
Distributions of the measured air temperatures T_ab_, T_in_ and the estimated panicle temperature T_p_ by IM^2^PACT, at the respective flowering 6 hours in the observation period, by box-plots. The “n.d.” means no data because of sensor malfunction. The mark “*” at the three seasons in IN, LK and MM means that the T_p_ could not be estimated because of no hourly data of solar radiation and wind speed. The mark “#” at the two seasons in IN and PH means that the T_p_ was estimated by using T_in_ and RH_in_ instead of missing data of T_ab_ and RHab, by solving the heat balance between the panicles and the air inside the canopy in IM^2^PACT.

#### 3.3.3 Sterility and panicle temperature at flowering hours

The sterility rate of IR64, N22 and IR52, were plotted against the thermal variables (Figure 8), where the thermal variables were averaged for 7 days from 50% heading day (days after heading, DAH, is 0 to 6) in the same way as in Figure 4. The correlation coefficient with the sterility rate tended to increase as using the thermal variable closer to the plant canopy. The sterility rate was most closely related to the thermal variable closest to the plant canopy, that is, the temperature of the panicles, which are sensing organs involved in heat induced spikelets sterility. Taking the average value of the T_p_ during the flowering time zone for 7 days including the heading date as a variable, the sterility rate of IR64 was expressed by the following equation, whose determination coefficient was 0.64.

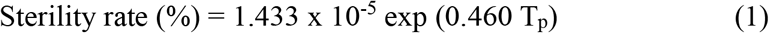

**Figure 8.**
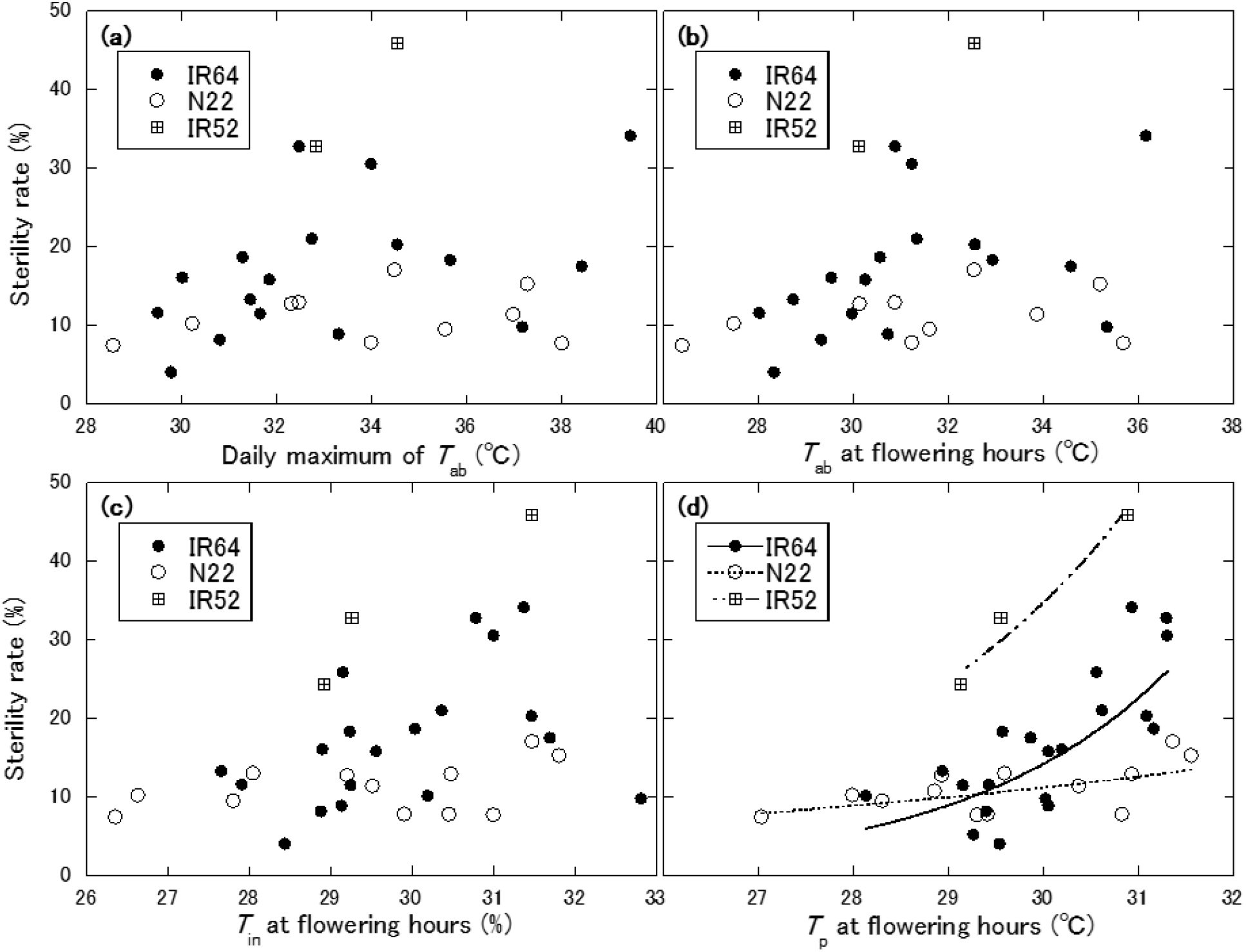
Relationship of the sterility rate of IR64, N22 and IR52 with thermal variables; (a) daily maximum T_ab_, (b) T_ab_ averaged in flowering 6 hours (c) T_in_ averaged in flowering 6 hours, and (d) T_p_ averaged in flowering 6 hours. All the thermal variables were averaged for 7 days from the respective heading day.

The sterility rate of N22 tended to increase slightly as the T_p_ increased, but it was almost constant. The relationship was expressed as below, and the determination coefficient was 0.39.

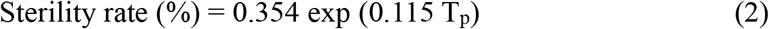

For IR52, the relationship was expressed as below.

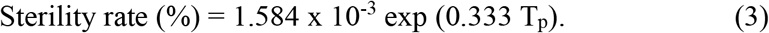

When the daily maximum T_ab_ or its average of flowering time zone are used as indicators, the difference between varieties in relation to the sterility rate is ambiguous. However, by using the T_p_ as the indicator, the difference in tolerance/susceptibility to HISS among varieties became apparent.

From the previous knowledge obtained at closed system experiments (e.g. Horie, 1993), the daily maximum air temperature of 35 °C has been often used as the temperature threshold for HISS. Bheemanahalli et al. (2016) identified the air temperature above the canopy of 33 °C as the critical threshold using 292 diverse indica cultivars grown in the field in the dry and wet seasons in India. It is suggested, however, that the threshold was lower, around 30 °C for IR64 for instance, when the T_p_ was used as the indicator.

## 4 DISCUSSION

### 4.1 The distribution of sterility among sites estimated by panicle temperature

This study showed the gap of micrometeorology among the nearby weather station, above the canopy and inside the canopy by the continuous and accurate measurement in the actual paddy fields in a wide variety of climates, which is the world’s first result that can be achieved by the multi-site monitoring network using the stand-alone force-ventilated measurement system to measure the air temperature and humidity, MINCER. Evapotranspiration by plant canopy and its evaporative cooling effect played an important role in the gap, so the climatic conditions of the site, especially the humidity environment, greatly affected the difference in temperature and humidity environment above and inside the canopy. The temperature of the panicles inside the canopy was also greatly affected by the humidity environment inside the canopy via its transpiration cooling effect, which further expanded the gap between the thermal environment above the canopy and the panicle temperature. Those gaps could be further complicated by the interaction of meteorological factors other than the humidity such as the solar radiation and wind speed. The correlation with the sterility rate of the common varieties was higher by using the temperature of the panicles, which are the flowering subjects, averaged during 6 hours related to the flowering event of each site, rather than by using the temperature above the canopy (Figure 8).

The distributions of possible sterility rate at all seasons shown in Figure 2 were estimated using the relational equations of (1) to (3) with the T_p_ averaged for the flowering hours (Figure 10), where the 7 days window of the flowering period (DAH = 0-6) was moved within 29 days to calculate the 7 days average of the T_p_ at flowering hours, which gave the possible sterility rate for the 23 cases from a season and a site at most.

The distribution of 7 days average of T_ab_ (for both its daily maximum and its flowering hours average) tended to be higher in the dry season than in the wet season in all tropical sites, and particularly high in the dry season in Senegal site. However, as approaching to the plant canopy and panicles as in T_in_ and T_p_, the temperature difference between the dry season and the wet season became smaller, and the T_p_ could be even higher in the wet season than in the dry season in Senegal site. In addition, the temperatures in wet climate like Benin site or in the temperate sites got closer to the level in the tropical dry seasons as approaching to the plant canopy and panicles. As a result, the sterility rate calculated by using the T_p_ was high in Benin, the Philippines (dry season) and China sites when compared by median of the distribution, and could also be high in the wet season in Senegal site, the dry season in Ghana site, and the USA site.

The humidity environment greatly affected the temperature gap between the air above the canopy and the panicle inside the canopy via the evaporative cooling of the canopy and panicles, which may change the global ‘hot spots’ in the light of the panicle temperature. Tian et al. (2010) reported that the panicle temperature was higher than the air temperature under humid and windless climatic condition, which induced the HISS in the field of Jianghan Basin in China despite the air temperature is not so high during flowering season. While in New South Wales in Australia, Matsui et al. (2014) observed that the panicle temperature was much lower than the air temperature above the canopy in paddy field because of the wind with very low humidity, which enabled low sterility even under the extreme hot climate. In the dry season of Senegal in this study, it is considered that the panicle temperature was low with the same heat balance structure as in New South Wales in Australia. Thus, by evaluating the sterility rate using the T_p_ as the indicator, there is a possibility that the risk map of HISS may be redrawn completely differently from ones often evaluated by using daily maximum T_ab_ or Tst.

### 4.2 Panicle temperature as an indicator for sterility

In order to draw the distribution of HISS in rice cultivation areas in the world, it is necessary to collect monitoring data of canopy micrometeorology over a longer period of time, and Figure 10 is not necessarily the exact distribution of the risk under the current climate, as they were estimated by using just one experimental period seemed to be typical weather condition chosen from each season and each site. Despite the fact, those results give us useful insights into the evaluation and adaptation research on the sterility risk.

One point is on the future projection of the heat sterility risk. Figures 9 and 10 showed that the HISS was high in the most humid not in the hottest environment. It was the same results as in Julia and Dingkuhn (2013) in which field experiments were conducted for monitoring of canopy microclimate and rice sterility in four contrasting environments including the dry seasons in Senegal and in the Philippines same as this study. This is because the T_p_ tended to rise due to suppression of evaporative cooling of canopy and panicles under the humid environment. Since such a heat balance structure is the same even after global warming, it is suggested that the risk of HISS in the future may increase in high humidity climates rather than in dry climates. It leads to the importance of the humidity accuracy as well as of the air temperature in climate scenarios and their spatial downscaling for future prediction of rice heat stress and production.

**Figure 9.**
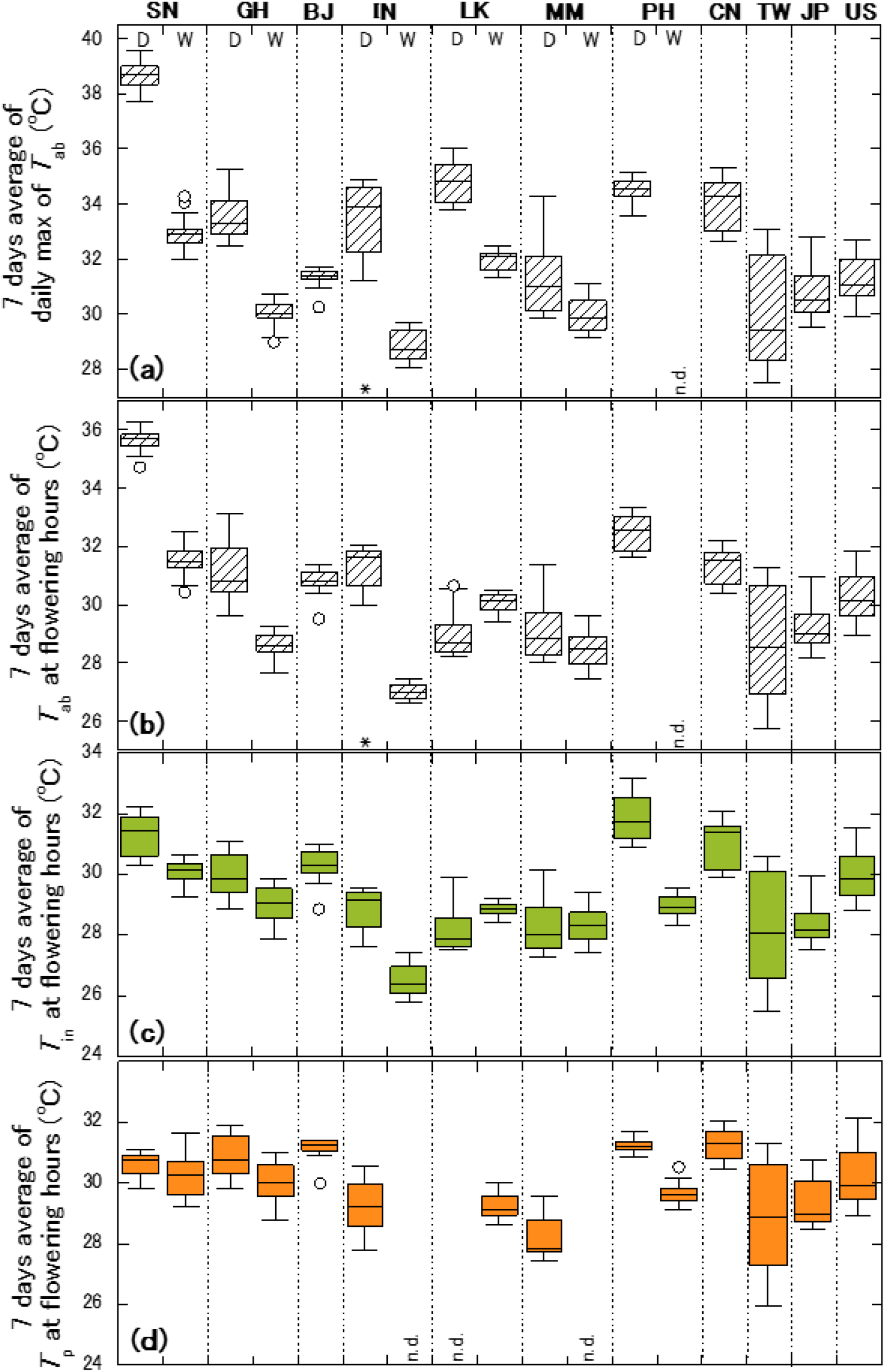
Distributions of 7 days average of thermal variables; (a) daily maximum T_ab_, (b) T_ab_ averaged in flowering 6 hours, (c) T_in_ averaged in flowering 6 hours, and (d) T_p_ averaged in flowering 6 hours, when the 7 days window of flowering period was moved within 29 days of observation period. “n.d” means that the T_p_ could not be estimated because of no hourly data of solar radiation and wind speed.

**Figure 10.**
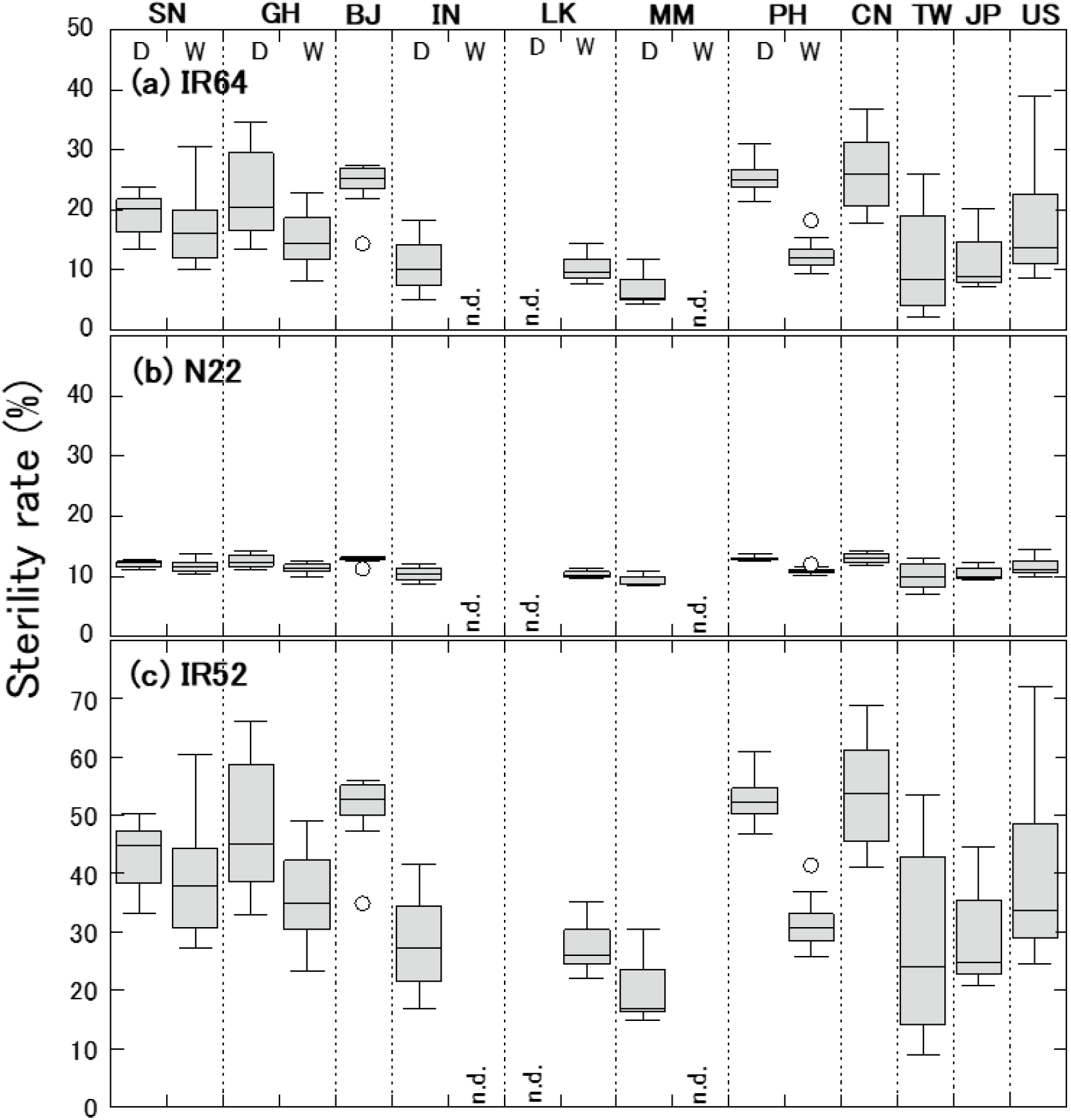
Distributions of sterility rate of (a) IR64, (b) N22 and (c) IR52, estimated by 7 days average of T_p_ at flowering, when the 7 days window of flowering period was moved within 29 days of observation period. “n.d” means no data because the T_p_ could not be estimated because of no hourly data of solar radiation and wind speed.

The evaluation of HISS by T_p_ can also contribute to the evaluation of the effectiveness of various adaptation measures. As adaptation measures, there are various options that account the ability of plants to tolerate, escape and avoid (Khan et al., 2019). As in Figure 8d, the genetically heat tolerant variety N22 had almost no increase in the sterility rate even when the T_p_ increased. On the other hand, the sterility rate of IR64, the moderately heat-susceptible variety, began to increase when the T_p_ exceeded 30 °C, and that of IR52, the heat susceptible variety, appeared to begin to increase at still lower T_p_. By using the T_p_ as the indicator, the temperature threshold value for each variety can be estimated and the sterility distribution can be quantitatively evaluated as shown in Figure 10. As a heat escape, it is considered as promising of the early morning flowering (EMF) trait that complete the flowering (spikelets opening) event before the temperature rises during the day, and varieties introducing EMF trait have been developed (Ishimaru et al., 2010; Hirabayashi et al., 2015). In recent years, MINCERnet has started the experiment cultivating the IR64 introduced of EMF (IR64-qEMF). It is possible to evaluate the effectiveness of EMF trait under a variety of climates by using the T_p_ during the respective flowering time as the indicator as in this study. As a heat avoidance, it is effective to directly cool the panicles by increasing the transpiration of the panicles themselves, in addition to cooling the panicles by the canopy transpiration. Fukuoka et al. (2012b) observed three time’s difference in the panicle transpirational conductance at flowering among 21 rice varieties in the field condition, which gave the estimation by IM^2^PACT of 3.5 °C difference in the T_p_ under the dry atmospheric condition. Using the T_p_ as an indicator, thus, makes it possible to set genetic improvement targets quantitatively.

Although it is certain that there is a high correlation between the sterility rate and the T_p_ at flowering hours, it has been generally difficult to evaluate the sterility rate using the T_p_ so far. One of reasons for this is the narrow temperature range the T_p_ can take, within which the response of the sterility rate to the T_p_ is steep. As shown in Fig. 8, for example in case of IR64, the temperature range was 10 °C or larger for the daily maximum T_ab_, about 10 °C for T_ab_ at flowering hours, and about 6 °C for T_in_ at flowering hours. The T_p_ at flowering hours, however, ranged only about 3 °C, and only 1 °C elevation in the T_p_ from 30 to 31 °C increased the sterility rate by nearly 10%. While, it is difficult to measure T_p_ continuously and to obtain the representative and accurate values from the measured T_p_ in the field because the actual T_p_ in the field can fluctuate every moment and have trend due to meteorological changes. That is the reason why it has been difficult to model the HISS in the field as a function of T_p_. The panicle temperature estimation by heat balance model such as IM^2^PACT is an effective means for developing the rice heat sterility model if there is the micrometeorological measurement data due to force-ventilation system accurate enough to apply to the heat balance equations. Alternatively instead of the T_p_, using the T_in_ as the indicator that can take larger temperature range than the T_p_, is seemed sufficiently effective as the second best measure for developing the HISS model, because the panicles are directly exposed to the air inside the canopy, and the T_in_ is more closely related to the sterility rate rather than the air temperature above the canopy, T_ab_, as shown in Fig. 8c. In that case, of course, the T_in_ must be measured by the force-ventilation system to obtain the accurate values inside the canopy in the field.

### 4.3 Micrometeorological gap to generate the uncertainties in the crop yield’s evaluation

Generally when experimentally investigating or modeling crop responses to the air temperature, it is common to measure the air temperature adjacent to but above the canopy or refer to the data of the nearby weather station as the variables that represent the thermal environment in the target field.

Regarding the temperature gap between the weather station and the farmland, Kuwagata et al. (2014) reported the case study in Japan, where the daily average of the air temperature measured at the meteorological station in urban site was 0.9 °C higher than that on the nearby farmland. In this study, however, even if the nearby weather station area was not urbanized, there was a gap between the T_st_ and T_ab_ mainly depending on the nearby weather station’s location (distance from the paddy field). Furthermore, there was a gap between T_ab_ and T_in_ mainly due to the evaporative cooling effect of plant canopy as in Figure 3. Those air temperature gaps are large enough to greatly affect the plant response predicted by using simple crop growth models based on cumulative temperature, or by using rice development models such as those of Horie (2019), Yoshida et al. (2010) and Bouman et al. (2001).

The most key problem is that the magnitude of the gap is not constant among sites and among seasons. Compared by the median of 24-hour values in Figure 2, for instance, in dry season in Senegal site, T_ab_ is 1.0 °C lower than T_st_ and T_in_ is still 2.4 °C lower than T_ab_, resulting in a gap of 3.4 °C between T_st_ and T_in_ actually sensed by plants. In dry season in Ghana site, the difference between T_st_ and T_ab_ was 1.1 °C and between T_ab_ and T_in_ was 1.7 °C, resulting in a gap of 2.8 °C between T_st_ and T_in_. While in wet sites such as Benin site and the temperate sites, the gap between T_st_ and T_in_ smaller than in dry sites in the range of 0.5 to 1.2 °C. And even in the same site, the gap between T_st_ and T_in_ may differ depending on whether the season is in the dry season or the wet season, such as in Ghana site.

Those gaps across diverse climates should generate great uncertainties in evaluating the impact of global crop production under climate change. To reduce such uncertainties, it should be effective to accumulate the reliable observation data inside the canopy, and to introduce and intervene a heat balance sub-model that can reproduce the thermal environment in the canopy. Data collected by MINCERnet hopefully contributes to crop model improvement activity such as AgMIP (Li et al., 2014) as a benchmark dataset, as well as the HISS evaluation by coping with the uncertainties due to the gap of the micrometeorology under global change.

## ACKNOWLEDGEMENTS

Our research was financially supported by the Global Environmental Research Coodination System from Ministry of the Environment of Japan, and JSPS KAKENHI Grant Number JP15H02650.

